# Aging increases the systemic molecular degree of inflammatory perturbation in patients with tuberculosis

**DOI:** 10.1101/2020.03.10.985697

**Authors:** Deivide Oliveira-de-Souza, Caian L. Vinhaes, María B. Arriaga, Nathella Pavan Kumar, Artur T. L. Queiroz, Kiyoshi F. Fukutani, Subash Babu, Bruno B. Andrade

## Abstract

Tuberculosis (TB) in a chronic infection that can affect individuals of all ages. The description of determinants of immunopathogenesis in TB is a field of tremendous interest due to the perspective of finding a reliable host-directed therapy to reduce disease burden. The association between specific biomarker profiles related to inflammation and the diverse clinical disease presentations in TB has been extensively studied in adults. However, relatively scarce data on profiling the inflammatory responses in pediatric TB are available. Here, we employed the molecular degree of perturbation (MDP) score adapted to plasma biomarkers in two distinct databanks from studies that examined either adults or children presenting with pulmonary or extrapulmonary disease. We used multidimensional statistical analyses to characterize the impact of age on the overall changes in the systemic inflammation profiles in subpopulation of TB patients. Our findings indicated that TB results in significant increases in MDP values, with the highest values being detected in adult patients. Furthermore, there were unique differences in the biomarker perturbation patterns and the overall degree of inflammation according to disease site and age. Importantly, the molecular degree of perturbation was not influenced by sex. Our results revealed that aging is an important determinant of the differences in quality and magnitude of systemic inflammatory perturbation in distinct clinical forms of TB.

## Introduction

Tuberculosis (TB) remains the leading cause of mortality worldwide due to a single agent [1]. *Mycobacterium Tuberculosis* (Mtb) is widely disseminated geographically and infects individuals of all ages, causing a wide spectrum of clinical manifestations associated with the host immunological status [2].

The majority of the studies exploring immunopathogenesis in TB is restricted to the adult population. On the other hand, TB pathophysiology remains poorly understood in children, especially in those under 5 years-old [3–5]. An important challenge in pediatric TB is the increased frequency of extrapulmonary presentations [6], which can be paucibacillary and thus associated with challenges in microbiologic test confirmation, resulting in delayed therapy implementation.

Inflammatory biomarkers in TB have been extensively studied and gained prominence due to the potential use as host-based blood tests [7]. Understanding the complexity of inflammatory milieu after TB infection in children is the key to expand the field and start development of new tools to aid the clinical management and contribute to non-sputum-based point of care test, with progress in diagnosis and clinical management.

Here, we used an adaptation of the molecular degree of perturbation (MDP), as previously published by us [8, 9] to assess the nuances of TB in pediatric patients compared to those observed in adult patient population. We found that the degree of inflammatory imbalance is associated with aging, suggesting that TB patients develop augmented capacity of promoting systemic inflammation with increasing age. Given that TB clinical presentation is a result of immunopathology, our results reinforce the idea that the distinct inflammatory profile in blood underlies the differences observed in TB disease presentation between adults and children.

## Materials and Methods

### Ethics statement

All clinical investigations were conducted according to the principles expressed in the Declaration of Helsinki. Written informed consent was obtained from all participants or their legally responsible guardians before enrolling into the sub-studies. The study was approved by the Institutional Review Board of the National Institute for Research in Tuberculosis, Chennai, India (NIRT; protocol numbers NCT01154959 and NCT00342017).

### Study design and participants

Active TB cases were recruited at the Government Stanley Medical Hospital, at TB clinics supported by the National Institute for Research in Tuberculosis and Childs Trust Hospital in Chennai, India. Detailed information on diagnosis of PTB in adults have been described previously in [8, 10] whereas the procedures used for diagnosis of PTB and EPTB in children have been reported in [11, 12]. Briefly, the diagnosis of PTB in adults and children was based on sputum smear and culture positivity. EPTB in adults was diagnosed on the basis of AFB staining and/or culture positivity of fine-needle aspiration biopsies of lymph nodes or pleural effusions in adult. The diagnosis of EPTB in children was made on clinical symptoms, physical examination and biopsies according to the site of clinical manifestation, such as fine-needle aspiration for the cases of TB lymphadenitis or cerebrospinal fluid analysis for TB meningitis as described in [12]. At the time of enrollment, all active TB cases had no record of prior TB disease or anti-TB treatment (ATT). The healthy control adults were asymptomatic with normal chest X-rays, negative TST (indurations < 5 mm in diameter) and QuantiFERON TB Gold-in-Tube enzyme-linked immunosorbent assay (Cellestis), as well as negative sputum smear or culture results as described in [8, 10]. Pediatric participants included in the heathy control group were asymptomatic who went to the hospital for routine vaccinations and tested negative in the QuantiFERON TB assay. All participants were BCG vaccinated and were HIV negative. Plasma samples were collected from a total of 152 adults and 54 children. Among adults, 97 were diagnosis of PTB, 35 of EPTB and 20 were healthy controls. Within children, 14 had PTB, 22 had EPTB and 18 were healthy controls. Adults with EPTB had TB lymphadenitis (n=24) or pleural TB (n=11) whereas children presented with spinal TB (n=14), TB lymphadenitis (n=6), and abdominal TB (n=2), which included peritonitis or tuberculomas. All participants were recruited in Chennai, India, as part of a large TB natural history study.

### Immunoassays

We evaluated a panel of 17 cytokines, tissue remodeling mediators and matrix metalloproteinases to examine molecular degree of inflammatory perturbation using different immunoassays. In the original study, biomarkers were measured in EDTA-treated plasma samples. Biomarkers included were cytokines, acute-phase proteins and tissue remodeling proteins. Bio-Plex multiplex ELISA cytokine assay system (R&D Systems) was employed to measure the cytokines analyzed. The list of cytokines included interleukin (IL)-1β, IL-10, IL-12p70, IL-17, interferon (IFN)-γ and tumor necrosis factor α (TNF-α). Moreover, IFN-α and IFN-β levels were quantified using the VeriKine serum ELISA kit (PBL Interferon Source). Plasma levels of vascular endothelial growth factor (VEGF) was measured using the Milliplex map kit system by Merck Millipore. Concentration of extracellular matrix metalloproteinases (MMPs)-1, 8 and 9, and tissue inhibitors of metalloproteinases (TIMPs)-1, 2, 3 and 4 were measured using Luminex technology (R&D Systems), according to the manufacturer’s protocols. Plasmatic Hemoxygenase 1 (HO-1) was measured by ELISA (Assay Designs).

### Data Analysis

Categorical data were presented as proportions and continuous data as medians and interquartile ranges (IQR). The Fisher’s exact test was used to compare categorical variables between study groups. Continuous variables were compared using the Mann-Whitney *U* test. P-values were adjusted for multiple measurements using Holm-Bonferroni’s method. Hierarchical cluster analyses (Ward’s method) of z-score normalized data were employed to depict the overall expression profile of indicated biomarkers in the study groups. Dendrograms represent Euclidean distance.

Profiles of correlations between biomarkers in different clinical groups were examined using network analysis of the Spearman correlation matrices (with 100X bootstrap). In indicated analyses, only correlations with significant adjusted P-value by the number of measurements (established cut-off was P-value <0.003) were included in the network visualization. In such analyses, markers that exhibited similar correlation profiles were clustered based on a modularity [13], which infers a sub-networks inside the of the correlation network profiles and depicted using Fruchterman Reingold (force-directed graph drawing)[14].

Sparse canonical correlation analysis (CCA) modeling was employed to assess whether combinations of circulating biomarkers could discriminate between subgroups of patients. The CCA model was chosen because many variables were studied. This model is able to perform dimensionality reduction for two co-dependent data sets (MDP biomarker profile and baseline characteristics profile, which were age and sex) simultaneously so that the discrimination of the clinical endpoints represents a combination of variables that are maximally correlated. Thus, trends of correlations between parameters in different clinical groups rather than their respective distribution within each group are the key components driving the discrimination outcome. In our CCA model, simplified and adapted from previously reported investigations of biomarkers for TB diagnosis [8, 15–18]. In the biomarker profile dataset, we included values of all the inflammatory marker variables. The diagnostic class prediction values obtained were calculated using receiver operator characteristics curve analysis. Probability of being molecularly perturbed according to increases in age was calculated using Kaplan-Meyer curves.

### Adaptation of the Molecular Degree of Perturbation to examine plasma concentrations of biomarkers

The plasma measurements of both datasets were normalized equally with a log2 transformation and the batch effect within the different study datasets was corrected using *Comba*t algorithm from SVA package [16]. The *ComBat* algorithm is a widely used method for adjusting batch effects in microarray and RNA-Seq data associated with technical variance effects. The molecular inflammatory perturbation is based molecular degree of perturbation (MDP) method used in the present study is an adaptation of the MDH described by Pankla et al. [19]. In the present study, instead of using gene expression values as in Prada-Medina et al. [20] we inputted plasma concentrations of a defined set of biomarkers pre-selected based on previously published studies from our group which investigated TB pathogenesis [15, 21]. Thus, herein, the average plasma concentration levels and standard deviation of a baseline reference group (healthy uninfected controls) were calculated for each biomarker. The MDP score of an individual biomarker was defined by the differences in concentration levels from the average of the biomarker in reference group divided by the reference standard deviation. Essentially, the MDP score represents the differences by number of standard deviations from the healthy control group. The formulas used to calculate MDP in the present study are shown below:

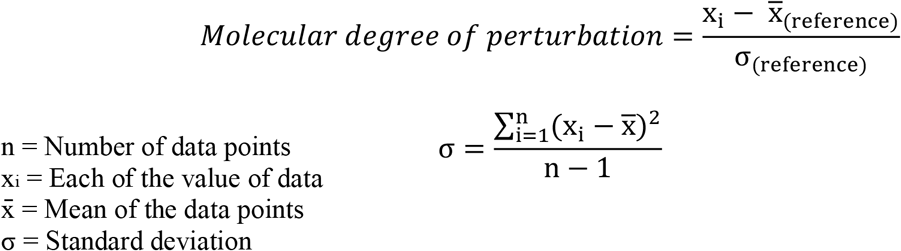

In this study, we applied the MDP scoring system using data on 17 biomarkers measured from two distinct groups of patients, adults and children with active TB and healthy uninfected controls. The MDP transformation was used as an approach to normalize data cross experiments resulting in datasets with markers distributed in a similar scale.

The MDP was filtered by the absolute MDP scores below 2 module and sum of all deviations accumulated MDP. To identify which sample was “perturbed”, we calculated the cutoff of the average MDP scores plus 2 standard deviations of the reference group and all values above this threshold was considered “perturbed”.

## Results

### Characteristics of study participants

Baseline characteristics of the adult and pediatric participants have been described elsewhere [12]. Adults with pulmonary TB and uninfected healthy controls were more frequently males than those with extrapulmonary TB (65.9% vs. 85% vs. 45.7%, respectively; P = 0.006). In addition, adults with PTB were on average older than healthy controls (median [IQR] in years: 38 [28-47] vs. 28.5 [26-35], respectively; P= 0.005) but had median age similar to the group of EPTB (**Table S1**). In the pediatric study, patients with PTB were similar to those presenting with EPTB or HC with regard age (median [IQR] in years: 6.5 [1.7-12.5] vs. 7 [3.7-13] vs. 10 [3-12], respectively; P = 0.684) and sex (P = 0.393), with and overall high frequency of male individuals (42.8% vs. 59.1% vs. 66.7%) (**Table S2**). We next compared the adults and children and found that there were no significant differences in sex distribution in comparisons between the HC, the PTB as well as the EPTB subgroups (**Table S3**).

### Increases in molecular degree perturbation of plasma cytokines and tissue remodeling mediators in adult and children with active tuberculosis

Plasma levels of 17 cytokines and tissue remodeling mediators were compared between pulmonary tuberculosis (PTB) and extrapulmonary tuberculosis (EPTB) and uninfected healthy controls (HC) in adult and children from India, separately (concentration values are described in **Table S4 and S5**). In adults, compared to HC, the PTB or EPTB groups exhibited higher levels of most parameters, except for IFN-β, TIMP-2, IL-12p70 and TIMP-4 which were not statistically different (**Table S4**). In the pediatric population, individuals with active TB (PTB or EPTB) exhibited on average higher levels of fewer markers (HO-1, MMP-1, MMP-8, TIMP-1 and TIMP-3) than controls (**Table S5**). These results suggested that TB in adults may lead to more significant changes in the concentration levels of plasma biomarkers than what we observed in children with this condition.

The data described above were originated from two different cohorts [10, 12]. In order to directly compare the groups from the distinct studies, we calculated the overall MDP score values according to our previous publication [8], that active TB was associated with a substantial increase in MDP scores compared to HC in both adults (PTB p<0.0001, EPTB p<0.0001; **Figure 1A**) and children (PTB p<0.001, EPTB p=0.002; **Figure 1B**). Adult patients with PTB exhibited higher MDP values than those with EPTB (p=0.0007; **Figure 1A**), whereas MDP values in PTB and EPTB were indistinguishable in children (p>0.999; **Figure 1B**). Interestingly, adult patients with either PTB or EPTB exhibited higher MDP values than those from the pediatric population (adults PTB vs. children PTB: p<0.0001; adults EPTB vs. children EPTB: p<0.001; **Figure 1C**) suggesting that the impact of Mtb on changes in the systemic molecular degree of perturbation is likely influenced by age. Reinforcing this idea, we found no differences in MDP values between adults and children from the HC group (p=0.46; **Figure 1D**).

**Figure 1:**
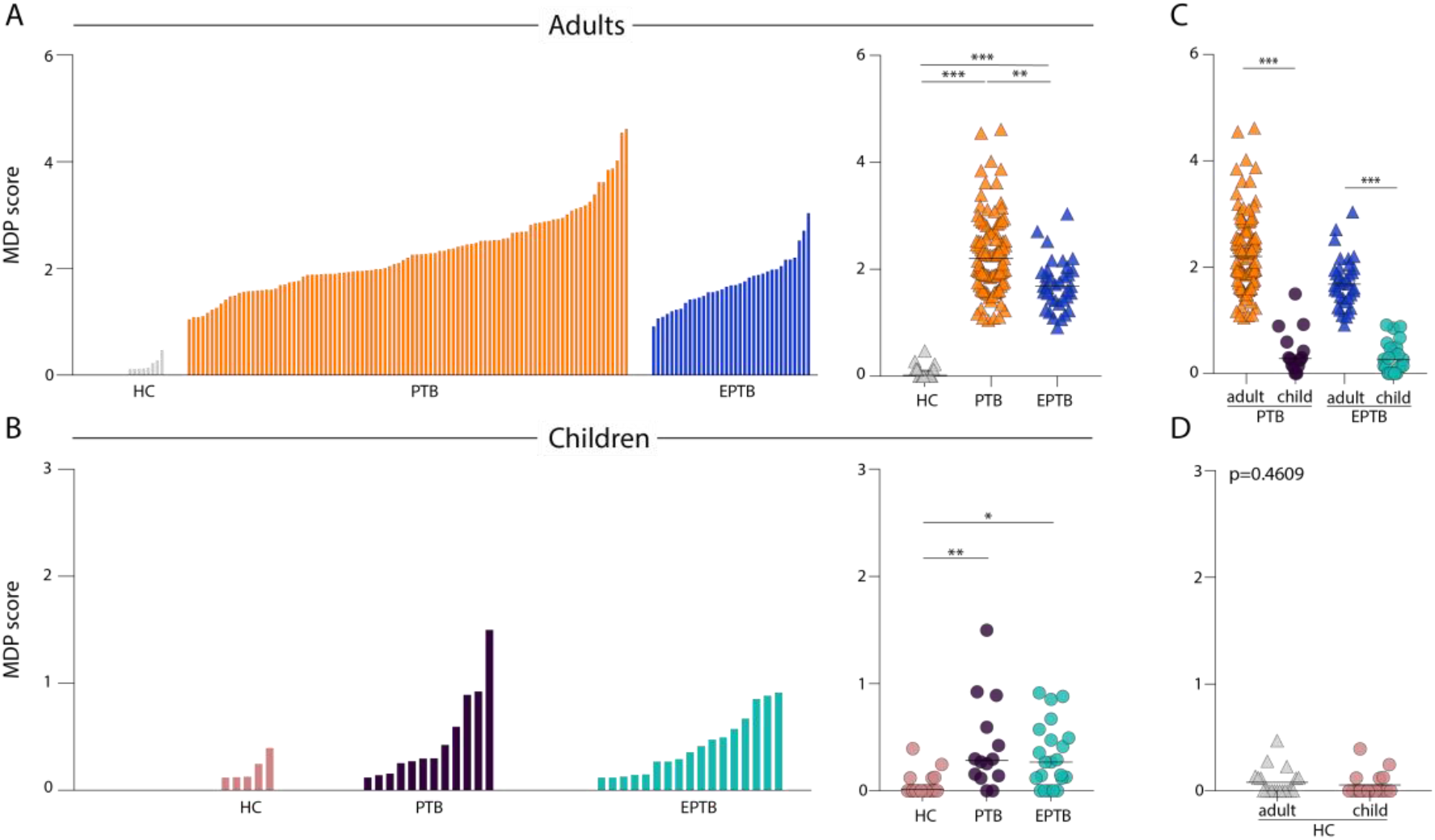
Adult and children with active tuberculosis exhibit substantial molecular degree of perturbation. **(A,B)** Left panels: Histograms show the single sample molecular degree of perturbation (MDP) score values relative to the healthy control group between adult and child (pulmonary TB: PTB, extrapulmonary TB: EPTB, healthy controls: HC). MDP values were calculated as described in Methods. The Kruskal Wallis test with Dunn’s multiple comparisons was used to compare MDP values between each clinical group. Right panels: Scatter plots of the summary data for each group are shown. MDP score values were compared between PTB or EPTB patients **(C)** or healthy controls **(D)** from Adult and Child. Lines in the scatter plots represent median values. Data were compared using the Mann-Whitney U test. *P<0.05; **P<0.01; ***P<0.0001.

### Plasma markers driving the overall molecular degree of perturbation in tuberculosis are distinct between adults and children

We examined the MDP expression values for each individual plasma cytokine and tissue remodeling mediators. An unsupervised hierarchical clustering was used to test whether the overall expression profile was associated with specific changes in active TB (PTB or EPTB) and HC in adults and children. We found that overall expression profile was very distinct between adults with active tuberculosis and HC. **(Figure 2A, left panel)**. Intriguingly, children showed heterogenous profile expression of the 17 markers between active TB and HC **(Figure 2B, left panel)**, without a clear separation in the cluster analysis. These findings indicate that the overall expression profile of the MDP values can be used to distinguish active TB from controls in adults but not in children. In addition, PTB could not be grouped separately from EPTB in both adults and children (**Figure 2A** and **Figure 2B**, **left panels**), indicating that active TB drives specific changes in MDP independent on disease site. Univariate analyses comparing the MDP values for each marker between HC and PTB or EPTB groups were showed in **Figure S1** and **Figure S2**.

**Figure 2.**
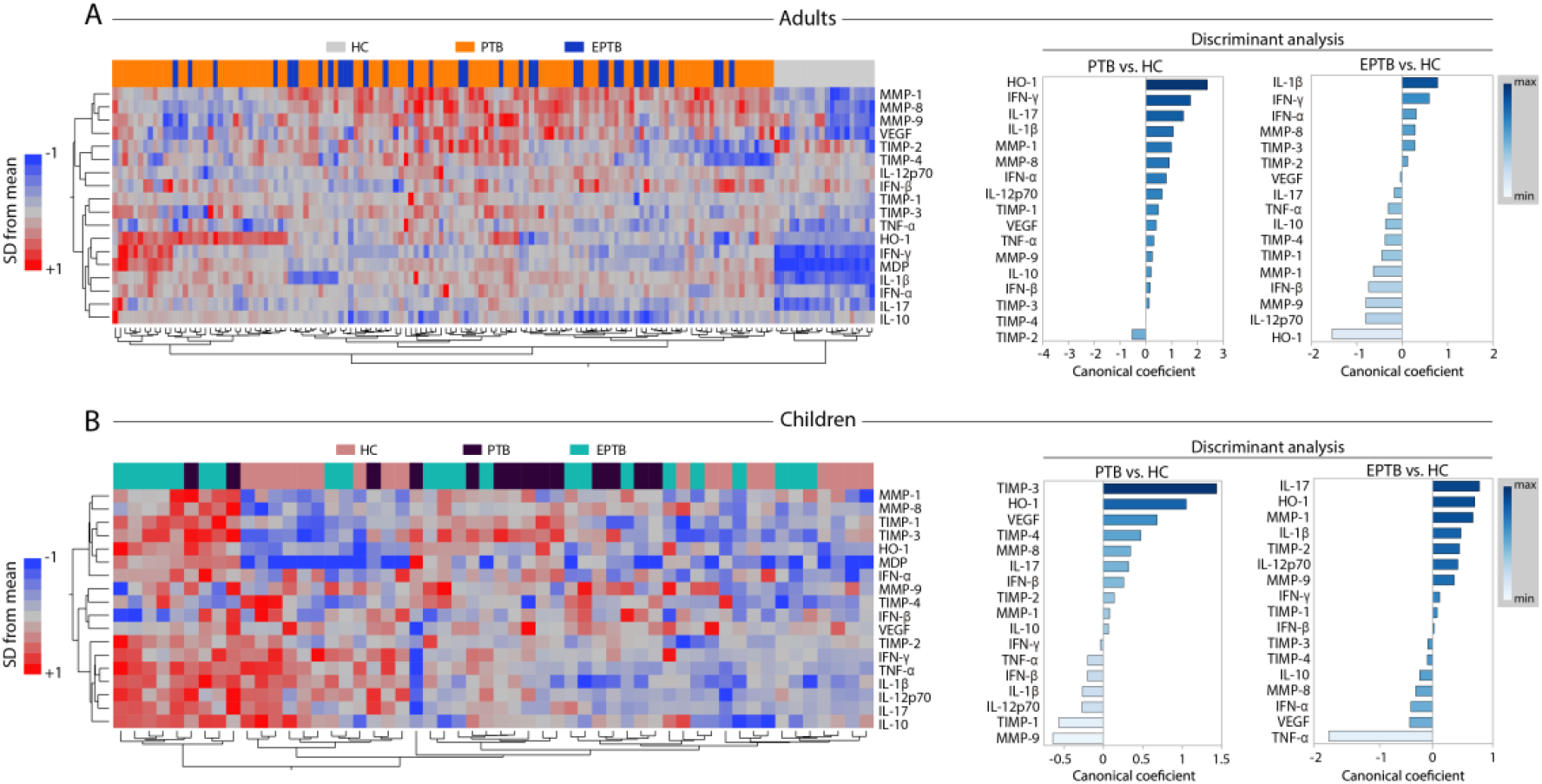
Plasma biomarkers driving the overall molecular degree of perturbation in pulmonary tuberculosis are distinct between Adult and Children patients. **(A,B)** Left panels: Unsupervised two-way hierarchical cluster analyses (Wards method with 100x bootstrap) using the MDP values for each individual markers measured in plasma from patients from both groups were employed to test if simultaneous assessment of such markers could group PTB or EPTB separately from healthy individuals. Dendrograms represent Euclidean distance. Right panels: A discriminant analysis model based on canonical correlation analyses was used to identify the markers which are driving the discrimination between the study groups. Number of patients per group: Adult HC: n = 20, Adult PTB: n = 97, Adult EPTB: n = 35, Child HC: n = 18, Child PTB: n = 14, Child EPTB: n = 22.

Furthermore, we employed a sparse canonical correlation analysis using MDP values [8] from all markers to identify which one contributes the most for the discrimination between PTB or EPTB and HC in both adults and children. Curiously, the markers with the strongest contributions for discrimination in adults with PTB vs. HC were HO-1, IFN-γ, IL-17, IL-1β and TIMP-2 whereas in children such markers were TIMP-3, HO-1, VEGF, TIMP-4 and MMP-9 **(Figure 2A-B right panels)**. The canonical model also indicated the markers that contributed the most for the discrimination EPTB vs. HC in adults were IL-1β, IFN-γ, IFN-α, MMP-8 and HO-1, whereas in children were IL-17, HO-1, MMP-1, IL-1β and TNF-α **(Figure 2A-B right panels)**. Of note, the MDP values of 3 out of 5 markers which were relevant to identify PTB in adults (HO-1, IFN-γ and IL-1β) were also part of TB signature observed in adults with EPTB. Moreover, in children, only 1 marker which was relevant to identify PTB (HO-1) was also relevant in EPTB. These findings indicate that in the context of Mtb infection, the plasma markers likely to be differentially perturbed according to disease site and age, except for HO-1.

### Network correlation profiles of molecular degree of perturbation in active TB are distinct between adults and children

To understand the nuances between molecular degree of perturbation of individual markers and their direct effect on overall MDP values, we employed network analysis based on Spearman correlation matrices, as previously described [8]. Using this approach, we found that the presence of pulmonary infection in adults was associated a greater number of correlations when compared with those that developed EPTB **(Figure 3A)**.The group of adults with PTB was marked by several positive correlations, highlighting that the degree of perturbation in HO-1, IFN-α, IFN-γ, IL-10, IL-17, IL-1β and TIMP-1 markers was directly associated with the overall MDP (**Figure 3A, left panel)**. In addition, perturbation of MMP-1 was inversely correlated with the overall MDP values in this clinical group. Furthermore, the top nodes exhibiting the highest number of significant correlations in the network of adults with PTB were MDP followed by MMP-1, IL-10, HO-1 and MMP-8. Importantly, the correlation profile found in the adult with EPTB was distinct from PTB (**Figure 3A)**. Indeed, only the degree of perturbation of HO-1 and IFN-γ were directly associated with the overall MDP values in the EPTB group. Node analysis of EPTB network demonstrated that MMP-8, IFN-γ, overall MDP and HO-1 and were the most highly connected parameters (**Figure 3A, right panel)**.

**Figure 3:**
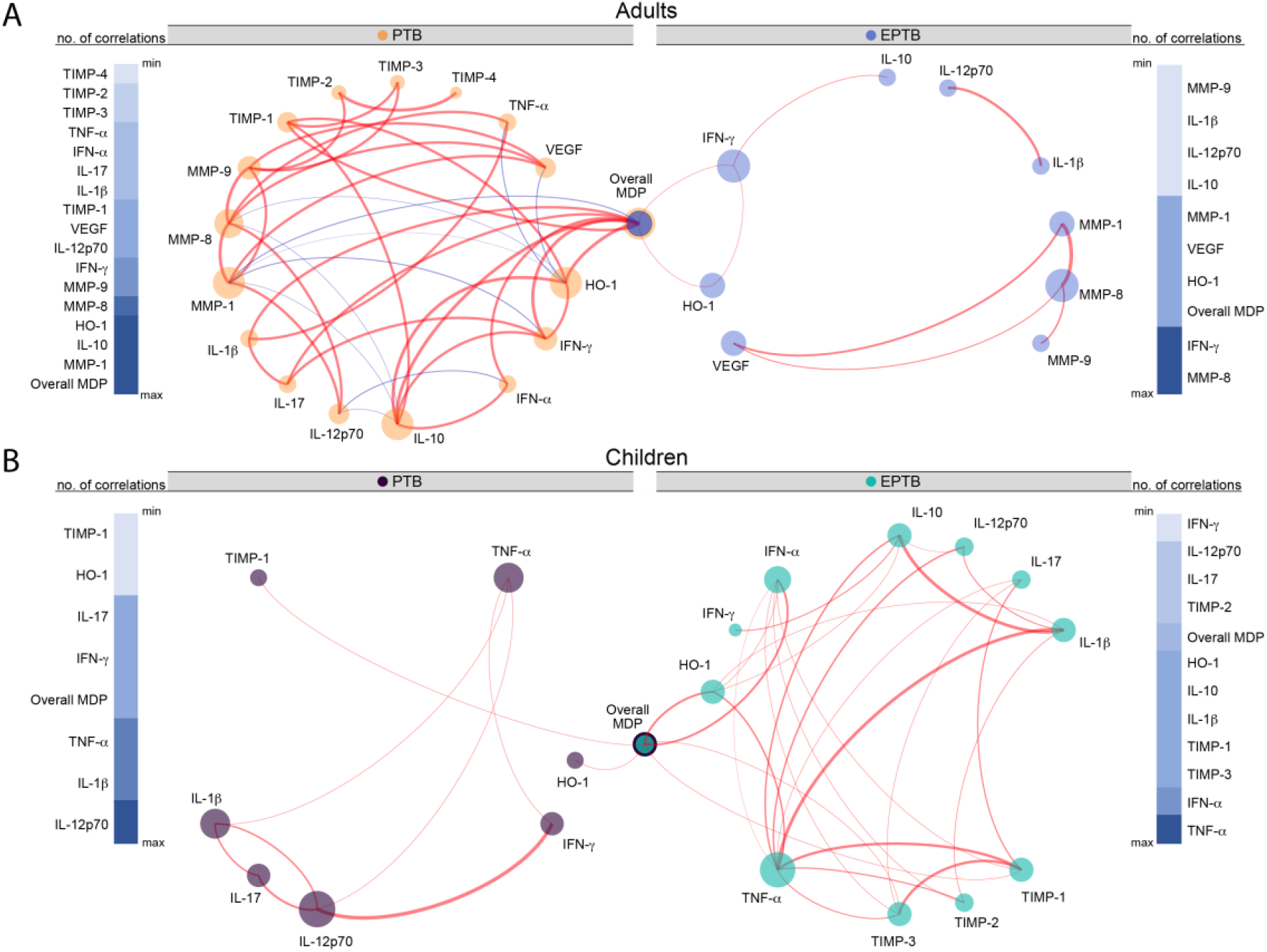
Network analysis of the MDP matrices in the study groups. **(A,B)** Spearman correlation matrices of the biomarker expression levels in each study group were built and Circos plots were used to illustrate the correlation networks. Each circle represents a different plasma parameter. The size of each circle is proportional to the number of significant correlations. P-values were adjusted for multiple measurements using Holm-Bonferroni’s method and the connecting lines represent statistically significant correlations (p<0.00277). Red connecting lines represent positive correlations while blue lines infer negative correlations. Color intensity is proportional to the strength of correlation (rho value). Node analysis was used to illustrate the number of significant correlations per marker. Markers were grouped according to the number of connections from minimum to maximum numbers detected.

When the network analysis was extended to the pediatric population, we observed a decreased number of statistically significant correlations in pulmonary infection when compared with extrapulmonary TB **(Figure 3B)**. Interestingly, we found that Mtb infection was associated with marked absence of negative correlations in children. Node analysis of the PTB network indicated that IL-12p70, IL-1β and TNF-α were the most highly connected markers **(Figure 3B, left panel)**. Curiously, the overall MDP values, which were highly connected in the networks from adults, were statistically correlated only with perturbation of HO-1 and TIMP-1. Children with extrapulmonary TB had TNF-α, followed by IFN-α, TIMP-3 and TIMP-1 as the most relevant nodes **(Figure 3B, right panel)**. The degree of perturbation of HO-1, IFN-α, TIMP-1 and TIMP-3 was directly associated with the overall MDP values in children with EPTB. Furthermore, in children, there was a lack of correlations between the MDP values of markers described to be important in TB pathogenesis such as IFN-γ, IL-1β and TNF-α in the networks. These findings argue that age influences the specific molecular markers that drive the overall molecular perturbation in active TB in adults and children.

### Age directly influences the overall inflammatory perturbation profile in active tuberculosis independent of sex

The results described above suggested that age was associated with the molecular degree of inflammatory perturbation. To directly test this hypothesis, we grouped each individual based on age and performed an exploratory investigation using unsupervised hierarchical cluster analysis with z-score normalized values of the MDP calculated for marker. We found that PTB did not exhibit a distinct biomarker MDP profile compared to EPTB independent on age **(Figure 4A, left panel**). Of note, MDP values detected for many markers were relatively lower in children independent on the TB clinical presentation, except for TIMP-1, TIMP-2, TIMP-3 and TIMP-4, which tended to be higher in those who were younger (**Figure 4A, left panel**). We next tested direct correlations between age and the individual MDP values of each marker. Spearman correlation analysis revealed that the degree of perturbation of IFN-γ, IL-1β, TNF-α, IFN-α, HO-1, IL-17, MMP-1, MMP-8 and VEGF was directly correlated whereas TIMP-2 values were inversely correlated with age **(Figure 4A, right panel)**.

**Figure 4:**
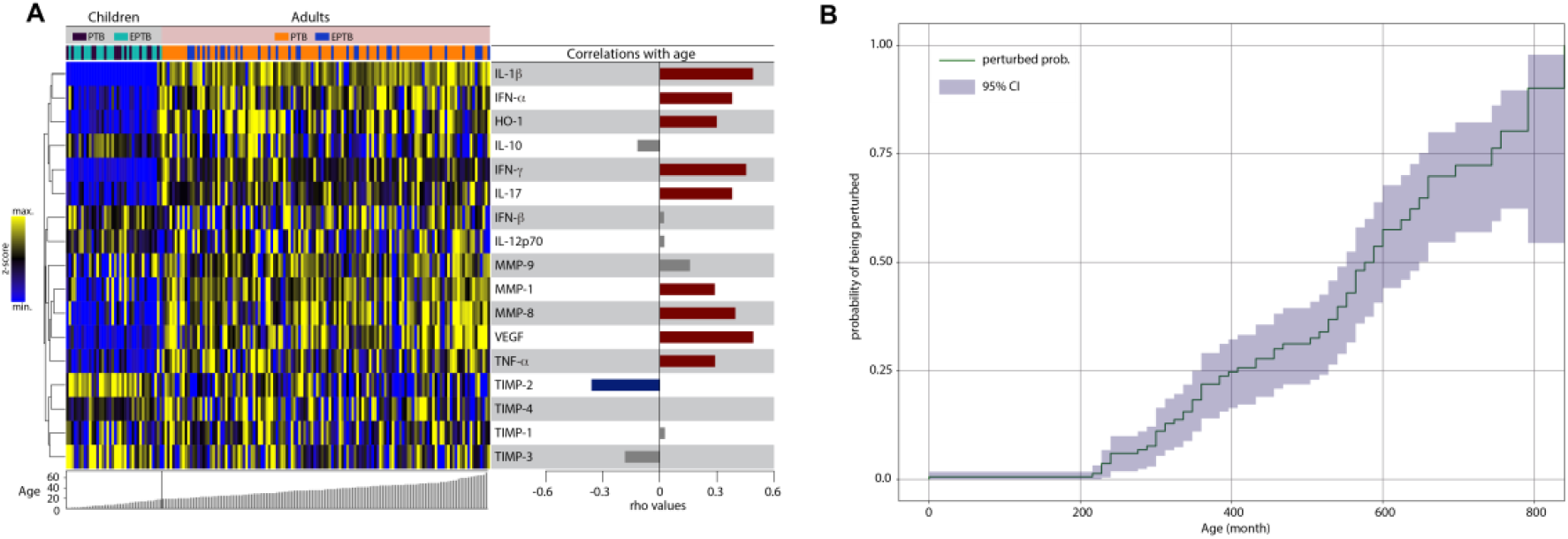
Associations between molecular degree inflammatory perturbation and age in TB patients. **(A)** Molecular degree of perturbation was assessed in samples from Adult and Child patients with tuberculosis. Data were z-score normalized. A hierarchical cluster analysis was employed to test whether the overall expression profile of the biomarkers could separate the study groups. Dendrograms represent Euclidean distance. Each individual was grouped based on age. The right panel shows Spearman correlation coefficient values of relationships between the indicated parameters and age. **(B)** Perturbed probability according to the age. Spearman correlation rank was compared used Steger Method.

To determine association between age and probability of being molecularly perturbed (see Methods for definition) in the entire population, we used a model adapted from the Kaplan-Meier survival curve (**Figure 4B)**. This approach revealed that increase in age was directly associated with higher probability of overall molecular perturbation. Indeed, the overall MDP score values were positive correlated with age in all the clinical groups evaluated **(Figure S3)**. Finally, we examined the influence of sex on the association between age and inflammatory perturbation. Overall MDP values were not different between male and female individuals stratified in the distinct clinical groups (**Figure 5A**). Furthermore, using the Kaplan-Meier survival curve test, we found that, in general, there was no different in the curves of female and male participants (p=0.29, **Figure 5A)**. Spearman correlation analysis demonstrated that MDP values of most of markers were positively correlated with age in both male and female participants **(Figure 5B)**. TIMP-2 was the only markers with a negative correlation in females. These findings suggested that sex does not influence the higher probability of inflammatory perturbation with aging in TB.

**Figure 5:**
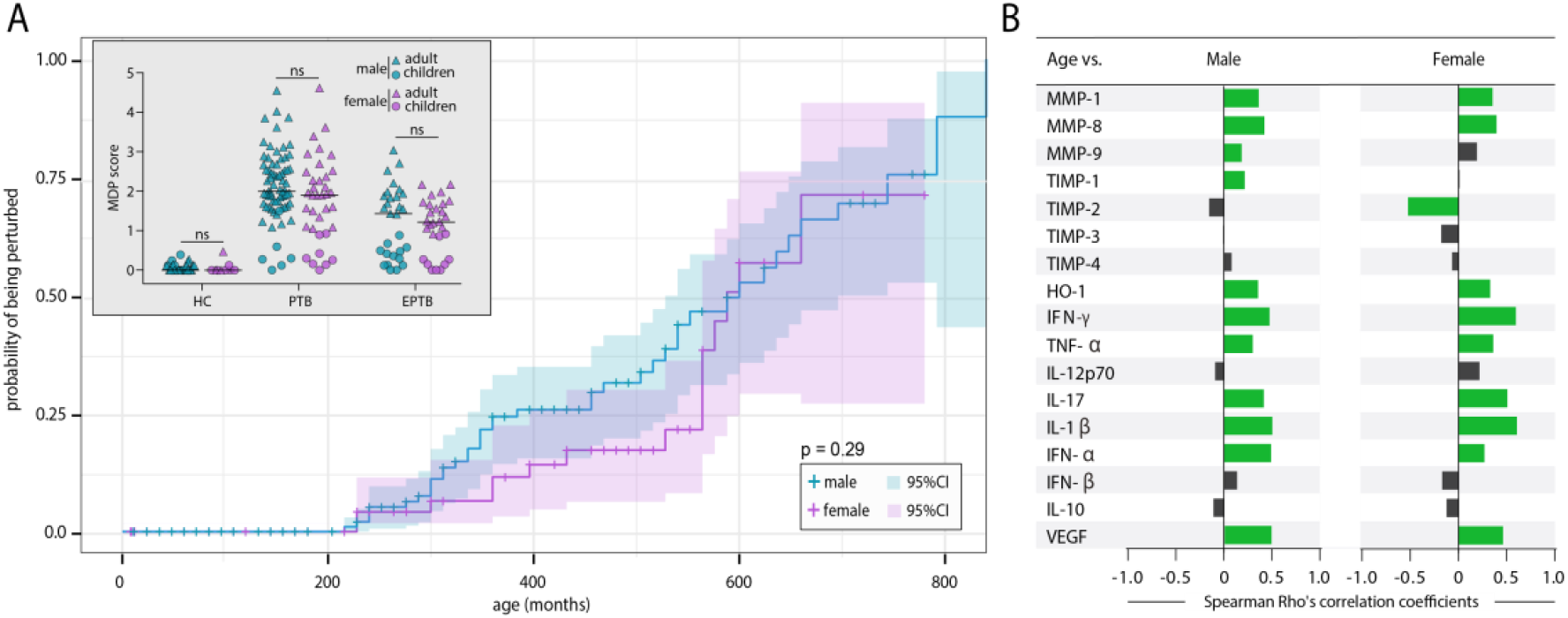
Molecular degree of perturbation is gender independent in patients with active tuberculosis. **(A)** Perturbed probability according to the age. Spearman correlation rank was compared used Steger Method. **(B)** Spearman correlation analysis was used to test association between age and gender. Bars represent the Spearman rank (rho) values. Colored bars indicate statistically significant correlation (p < 0.05) after adjustment for multiple measurement.

## Discussion

Mechanisms of disease pathogenesis in TB have been extensively studied over the years, but despite that, the immunopathology of this infection in pediatric populations remains poorly understood. Importantly, Mtb infection is one of the main causes of childhood morbidity and mortality worldwide [11, 22, 23], and the field needs elucidation of the determinants of immunopathogenesis. In the present study, we used an adaption of molecular degree of perturbation [8, 9] to estimate the level and quality of systemic inflammation in patients with active TB (PTB and EPTB) according to age. Our findings indicated that there are important discrepancies in the MDP values between adults and children with active TB, with adults exhibiting higher values, whereas the molecular perturbation was similar among individuals from the healthy control groups independent of age. Thus, while the systemic inflammatory profile is similar between adults and children without TB, it becomes very distinct in patients with active disease. Although we have not directly tested potential influence of maturation of immune system in the results, it is possible that the capacity to promote significant systemic inflammation in the context of TB may be affected by this process.

Mycobacterial infection is known to cause profound stimulation of both innate and adaptive immune response, in vitro and in vivo models [24, 25]. Our exploratory analysis characterized the systemic inflammatory response and indicated that Mtb infection (pulmonary and extrapulmonary) was associated with overall increases in the MDP values in both adults and children. However, in either PTB or EPTB groups, adult patients exhibited augmented MDP values compared to pediatric patients. This finding reinforces the idea that in the context of TB, adults are more prone towards presenting with a higher degree of inflammation than children. We expanded these analyses to show the degree of perturbation of each individual biomarker and found that the in adults, individuals with active TB exhibit a very distinct profile compared to those without. Nevertheless, in the pediatric population, there was no clear combined biomarker profile that could distinguish TB from controls. Hence, there are specific changes in the biomarker MDP profile that are age dependent. Of note, using discriminant analyses based on a canonical model, we identified that the top markers responsible for the discrimination between the clinical groups differed between adults and children. In adults, HO-1, IFN-γ and IL-1β, markers which have been associated with TB pathogenesis [26–28] were the top markers that contributed to discrimination between the disease groups (both PTB and EPTB) and controls[15, 26, 29] whereas in children, TIMP-3, IL-17 and HO-1 were the markers that most contributed for discrimination between active TB patients and uninfected controls. Of note, important finding of this analytical approach was that the molecular perturbation of HO-1 could discriminate pulmonary and extrapulmonary TB from healthy controls in both adult and children populations. HO-1 is important marker of oxidative stress response, highly expressed in the lungs [10, 30], with a critical role in cytoprotection [31]. These observations made us hypothesize HO-1 may be important in TB pathogenesis regardless of age. Additional studies in more diverse populations are warranted to test this idea.

The inflammatory process results from an intricate relationship between factors from the host and pathogen, and can be evaluated using network analysis [9, 32, 33]. Using this approach, we showed important differences in the correlation profiles between MDP values from each individual biomarker and the overall MDP values in adults and children with active TB. Pulmonary infection with Mtb in adults led to a coordinated inflammatory burst, which was read by detection of many statistically significant correlations between individual markers and overall MDP values. On the converse, in children presenting with PTB the augmented inflammation seemed less coordinated, meaning increases in molecular perturbation of a given marker were not followed by simultaneous increases of other markers or the overall MDP values. This suggests that pediatric PTB patients may have a decreased ability to mount a coordinated response. It is possible to hypothesize that such uncoupling of inflammatory activity may be a consequence of immune immaturity that leads to dissemination of bacilli in children, as previously showed [6]. Curiously, in children with EPTB, the networks were more complex, indicating a higher number of connections. Such phenomenon could be consequence of extrapulmonary tissue damage, and maybe argue that despite the lower capacity to mount and sustain a coordinated response in pulmonary infection, EPTB in children is marked by a more balanced interplay between innate and adaptive immune response [24]. Moreover, in adults with EPTB, the complexity of the inflammatory network was reduced, in agreement with the previous published evidence of probably less organized and an unfettered immune activation in adults with EPTB [9]. The determinants of the differences in correlations between concentrations of mediators of inflammation between distinct clinical forms of TB in adults and children are still unknown.

An interesting result reported here was the association between increases in age and augmentation of the overall molecular degree of perturbation in both PTB and EPTB patients. Indeed, our results demonstrated that increase in age leads to rise in probability of a TB patient becoming more perturbed, independent on disease site (PTB or EPTB). Importantly, we showed that degree of perturbation of many individual plasma markers correlated with age, reinforcing the idea that the capacity to induce systemic inflammation is proportional to age. Furthermore, our analysis showed that sex did not influence the probability of increase in MDP values, arguing that the potential effect of sex-related hormones on systemic inflammation may be superposed by the effect of aging in TB patients.

Our study has limitations. We were unable to test association between MDP and bacillary loads due to lack of data from the pediatric population. It is possible that higher MDP values detected in adults may be a consequence of the increased mycobacterial infection loads. In addition, the sample size of the group of children was relatively small. Regardless, the extensive exploratory analyses performed revealed unique relationships between age and the systemic degree of inflammation. The results presented here shed light into the impact of aging on the systemic immune activation during TB.

## Acknowledgments

The authors acknowledge study participants. This project was supported by the Intramural Research Program of the NIAID to S.B. and N.P.K. This study was also financed in part by Coordenação de Aperfeiçoamento de Pessoal de Nível Superior (CAPES) (Finance Code 001). The work of B.B.A. was supported by grants from the NIH (U01AI115940, R01AI069923-08, R01AI20790-02), by Intramural Program of Fundação Oswaldo Cruz and by the Brazilian National Council for Scientific and Technological Development (CNPq). D.O.S. and M.B.A. receive fellowships from the Fundação de Amparo à Pesquisa da Bahia (FAPESB). C.L.V. is a research fellow and K.F.F. is a postdoctoral fellow from CNPq.

## Author contributions

B.B.A. designed the study and mentored the work; D.O.S., C.L.V., M.B.A., N.P.K. performed the experiments and data collection; B.B.A., K.F.F., A.T.L.Q., M.B.A, D.O.S., C.L.V. performed data analyses; S.B. coordinated the clinical study, provided reagents for the immunoassays and helped with data interpretation; B.B.A., D.O.S., C.L.V., S.B. wrote the manuscript. All authors have read and approved the final version of the manuscript.

## Data availability statement

The datasets generated during and/or analyzed during the current study are available from the corresponding author on reasonable request.

## Supplemental Tables and Figures

**Table S1.** Characteristics of adult participants.

| Characteristic | Healthy controls | PTB | EPTB | P-value |
| --- | --- | --- | --- | --- |
| N | 20 | 97 | 35 |  |
| Age – y | 28.5 (26-35) | 38 (28-47) | 35 (25-45) | 0.0452 |
| Male – no. (%) | 17 (85) | 64 (65.9) | 16 (45.7) | 0.0062 |

Data represent medians and interquartile ranges (age) and frequencies (male sex). The Kruskal-Wallis test was used to compare distributions of age while the Pearson's chi-square test was used to compare frequencies.

**Table S2.** Characteristics of pediatric participants.

| Characteristic | Healthy controls | PTB | EPTB | P-value |
| --- | --- | --- | --- | --- |
| N | 18 | 14 | 22 |  |
| Age – y | 10 (3-12) | 6.5 (1.7-12.5) | 7 (3.7-13) | 0.6841 |
| Male – no. (%) | 12 (66.7) | 6 (42.8) | 13 (59.1) | 0.3928 |

Data represent medians and interquartile ranges (age) and frequencies (male sex). The Kruskal-Wallis test was used to compare distributions of age while the Pearson’s chi-square test was used to compare frequencies.

**Table S3.** Sex distribution in study population.

| Group | Sex | Adult | Children | P-value |
| --- | --- | --- | --- | --- |
| HC | Male | 17 | 12 | 0.2603 |
|  | Female | 3 | 6 |  |
| PTB | Male | 64 | 6 | 0.1372 |
|  | Female | 33 | 8 |  |
| EPTB | Male | 16 | 13 | 0.4173 |
|  | Female | 19 | 9 |  |

Data represent number of individuals in each group/category. The Fisher’s exact test was used to compare frequencies.

**Table S4.** Distribution of the plasma concentrations of the mediators of inflammation in adult participants.

| Parameter | Unit | Healthy controls | PTB | EPTB | P-value |
| --- | --- | --- | --- | --- | --- |
| N |  | 20 | 97 | 35 |  |
| Heme Oxygenase-1 | ng/ml | 1.5<br>(1.3-1.6) | 5.6<br>(3.2-11.6) | 3.8<br>(2.3-6.5) | **<0.0001** |
| IFN-α | pg/mL | 3.2<br>(1-4.3) | 11.6<br>(7.9-17.3) | 8.3<br>(4.8-12.1) | **<0.0001** |
| IFN-β | pg/mL | 4.4<br>(2.2-5.5) | 2.9<br>(1.3-7.1) | 3.4<br>(1.5-5.9) | 0.8231 |
| IFN-γ | pg/mL | 20.6<br>(15-19.5) | 33.9<br>(29.8-35.3) | 29.8<br>(27.8-36.0) | **<0.0001** |
| IL-10 | pg/mL | 18.4<br>(16.43-20.85) | 21.0<br>(15.93-29.05) | 14.8<br>(10.3-21.0) | **0.0014** |
| IL-12p70 | pg/mL | 2.3<br>(1.6-3.3) | 2.2<br>(0.9-5.3) | 2.9<br>(1.5-6.9) | 0.2078 |
| IL-17 | pg/mL | 16.3<br>(43.1-493.1) | 27.1<br>(25.85-35.32) | 26.5<br>(23.2-30.0) | **<0.0001** |
| IL-1β | pg/mL | 2.8<br>(2.5-3.4) | 15.9<br>(12.9-20.1) | 13.1<br>(11.7-15.9) | **<0.0001** |
| MMP-1 | ng/mL | 0.07<br>(0.04-0.32) | 3.4<br>(1.9-5.3) | 2.9<br>(2.2-4.5) | **<0.0001** |
| MMP-8 | ng/mL | 13.9<br>(5.1-32.5) | 122.4<br>(54.1-202.3) | 88.4<br>(75.6-141.9) | **<0.0001** |
| MMP-9 | ng/mL | 133.7<br>(53.7-331.5) | 384.2<br>(255.0-510.8) | 271.6<br>(214.7-329.6) | **<0.0001** |
| TIMP-1 | ng/mL | 151.4<br>(136.0-166.2) | 180.0<br>(161.7-199.7) | 169.6<br>(153.3-186.5) | **<0.0001** |
| TIMP-2 | ng/mL | 232.9<br>(219.2-255.9) | 218.9<br>(199.4-238.7) | 223.0<br>(208.3-234.1) | 0.1001 |
| TIMP-3 | ng/mL | 12.4<br>(5.5-19.2) | 26.3<br>(15.9-43.9) | 19.8<br>(11.5-29.4) | **<0.0001** |
| TIMP-4 | ng/mL | 9.6<br>(8.4-10.8) | 10.3<br>(7.7-12.4) | 8.8<br>(7.5-10.9) | 0.2174 |
| TNF-α | pg/mL | 12.5<br>(10.2-17.5) | 21.3<br>(14.9-27.4) | 18.9<br>(15.5-23.9) | **0.0006** |
| VEGF | pg/mL | 26.0<br>(18.4-47.8) | 100.8<br>(57.9-165.5) | 71.5<br>(43.0-113.3) | **<0.0001** |

Data represent medians and interquartile ranges. The Kruskal-Wallis test was used to compare the distributions of the plasma mediators between the study groups. P-values in bold font are statistically significant.

**Table S5.**
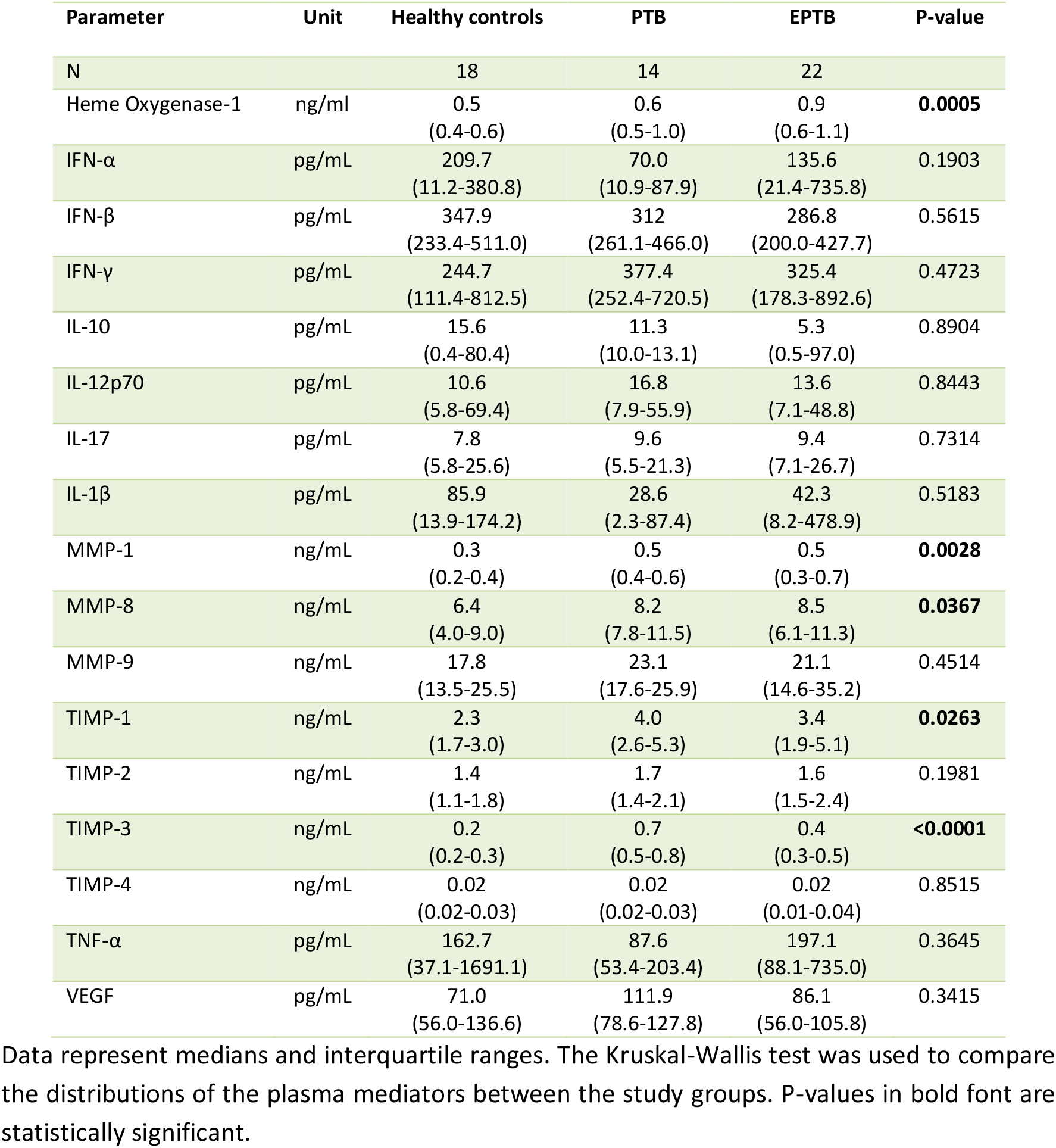
Distribution of the plasma concentrations of the mediators of inflammation in pediatric participants.

| Parameter | Unit | Healthy controls | PTB | EPTB | P-value |
| --- | --- | --- | --- | --- | --- |
| N |  | 18 | 14 | 22 |  |
| Heme Oxygenase-1 | ng/ml | 0.5<br>(0.4-0.6) | 0.6<br>(0.5-1.0) | 0.9<br>(0.6-1.1) | **0.0005** |
| IFN-α | pg/mL | 209.7<br>(11.2-380.8) | 70.0<br>(10.9-87.9) | 135.6<br>(21.4-735.8) | 0.1903 |
| IFN-β | pg/mL | 347.9<br>(233.4-511.0) | 312<br>(261.1-466.0) | 286.8<br>(200.0-427.7) | 0.5615 |
| IFN-γ | pg/mL | 244.7<br>(111.4-812.5) | 377.4<br>(252.4-720.5) | 325.4<br>(178.3-892.6) | 0.4723 |
| IL-10 | pg/mL | 15.6<br>(0.4-80.4) | 11.3<br>(10.0-13.1) | 5.3<br>(0.5-97.0) | 0.8904 |
| IL-12p70 | pg/mL | 10.6<br>(5.8-69.4) | 16.8<br>(7.9-55.9) | 13.6<br>(7.1-48.8) | 0.8443 |
| IL-17 | pg/mL | 7.8<br>(5.8-25.6) | 9.6<br>(5.5-21.3) | 9.4<br>(7.1-26.7) | 0.7314 |
| IL-1β | pg/mL | 85.9<br>(13.9-174.2) | 28.6<br>(2.3-87.4) | 42.3<br>(8.2-478.9) | 0.5183 |
| MMP-1 | ng/mL | 0.3<br>(0.2-0.4) | 0.5<br>(0.4-0.6) | 0.5<br>(0.3-0.7) | **0.0028** |
| MMP-8 | ng/mL | 6.4<br>(4.0-9.0) | 8.2<br>(7.8-11.5) | 8.5<br>(6.1-11.3) | **0.0367** |
| MMP-9 | ng/mL | 17.8<br>(13.5-25.5) | 23.1<br>(17.6-25.9) | 21.1<br>(14.6-35.2) | 0.4514 |
| TIMP-1 | ng/mL | 2.3<br>(1.7-3.0) | 4.0<br>(2.6-5.3) | 3.4<br>(1.9-5.1) | **0.0263** |
| TIMP-2 | ng/mL | 1.4<br>(1.1-1.8) | 1.7<br>(1.4-2.1) | 1.6<br>(1.5-2.4) | 0.1981 |
| TIMP-3 | ng/mL | 0.2<br>(0.2-0.3) | 0.7<br>(0.5-0.8) | 0.4<br>(0.3-0.5) | **<0.0001** |
| TIMP-4 | ng/mL | 0.02<br>(0.02-0.03) | 0.02<br>(0.02-0.03) | 0.02<br>(0.01-0.04) | 0.8515 |
| TNF-α | pg/mL | 162.7<br>(37.1-1691.1) | 87.6<br>(53.4-203.4) | 197.1<br>(88.1-735.0) | 0.3645 |
| VEGF | pg/mL | 71.0<br>(56.0-136.6) | 111.9<br>(78.6-127.8) | 86.1<br>(56.0-105.8) | 0.3415 |

Data represent medians and interquartile ranges. The Kruskal-Wallis test was used to compare the distributions of the plasma mediators between the study groups. P-values in bold font are statistically significant.

**Figure S1.**
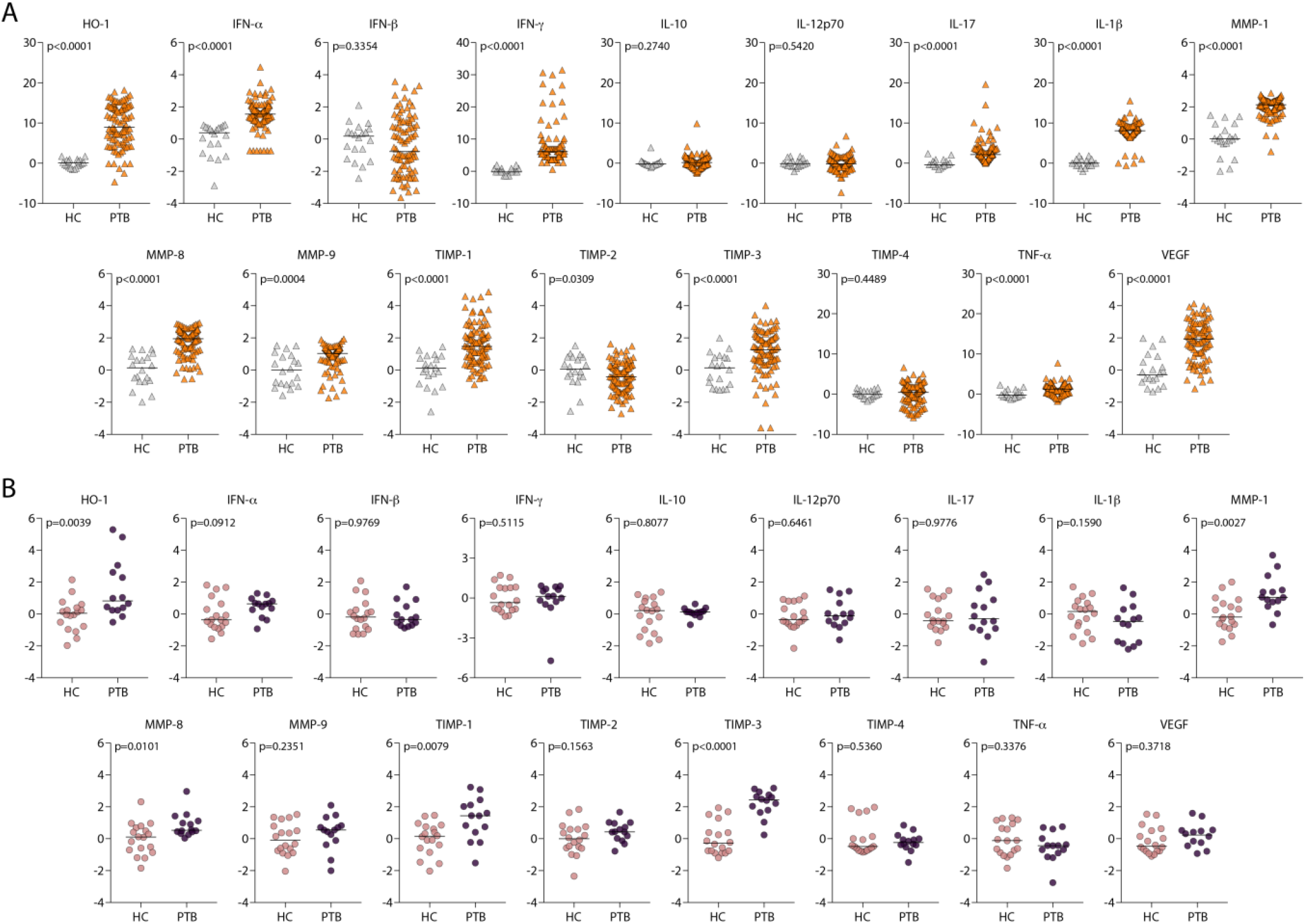
Differences in the inflammatory perturbation of each plasma biomarker between active PTB and healthy control groups in either adults or children. **(A,B)** Scatterplots of molecular degree of perturbation (MDP) of indicated biomarkers of TB patients stratified per the group (Adult HC, n=20 and PTB, n=97; Children HC, n=18 and PTB, n=14). MDP scores are shown in Y-axis from all the plots. Lines represent median values. Data were compared using the Mann–Whitney *U* test.

**Figure S2.**
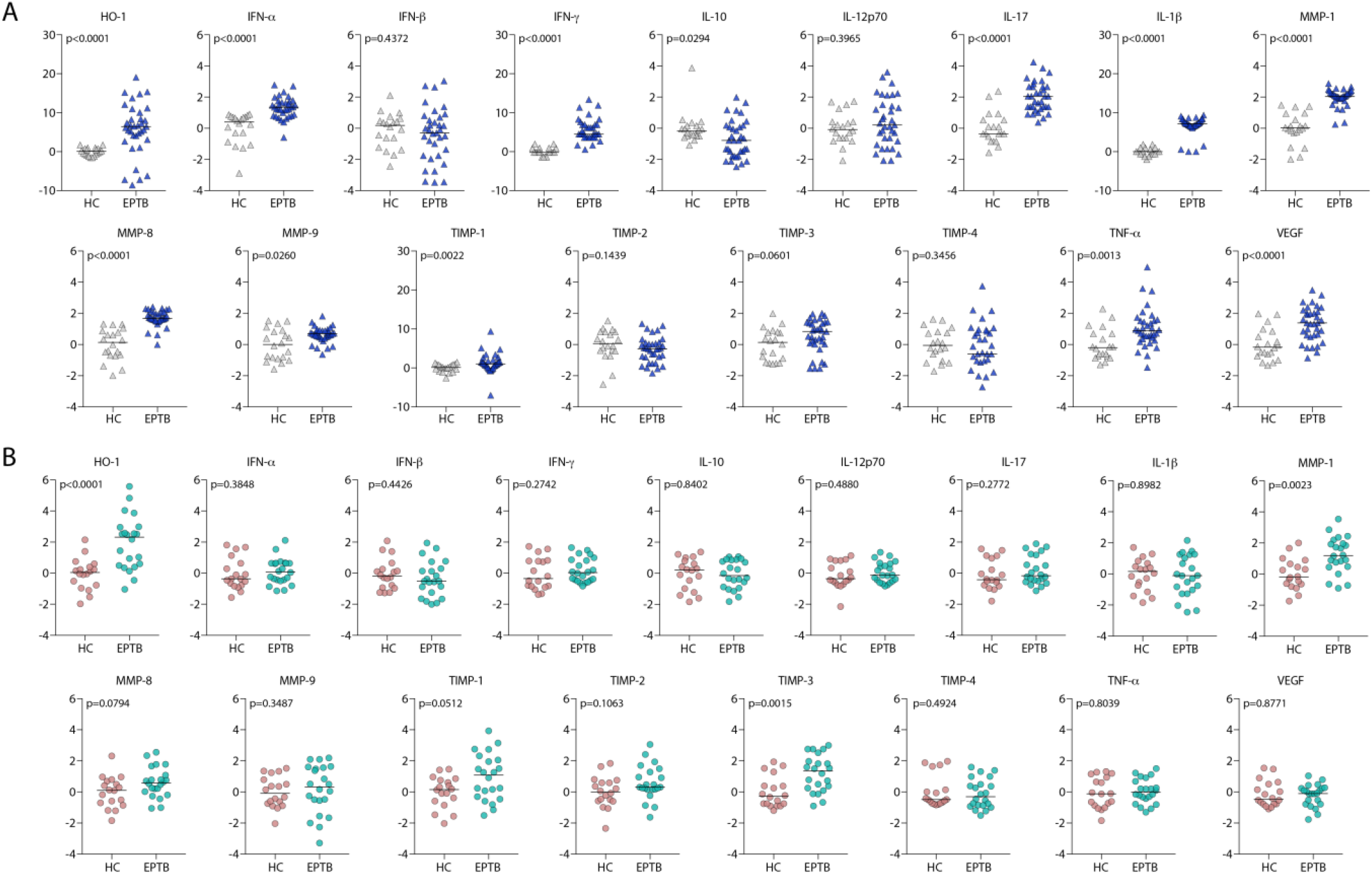
Differences in the inflammatory perturbation between adult or children with extrapulmonary tuberculosis compared to healthy controls. **(A,B)** Scatterplots of molecular degree of perturbation (MDP) of indicated biomarkers of TB patients stratified per the group (Adult HC, n=20 and EPTB, n=35; Children HC, n=18 and EPTB, n=22). MDP scores are shown in Y-axis from all the plots. Lines represent median values. Data were compared using the Mann–Whitney *U* test.

**Figure S3.**
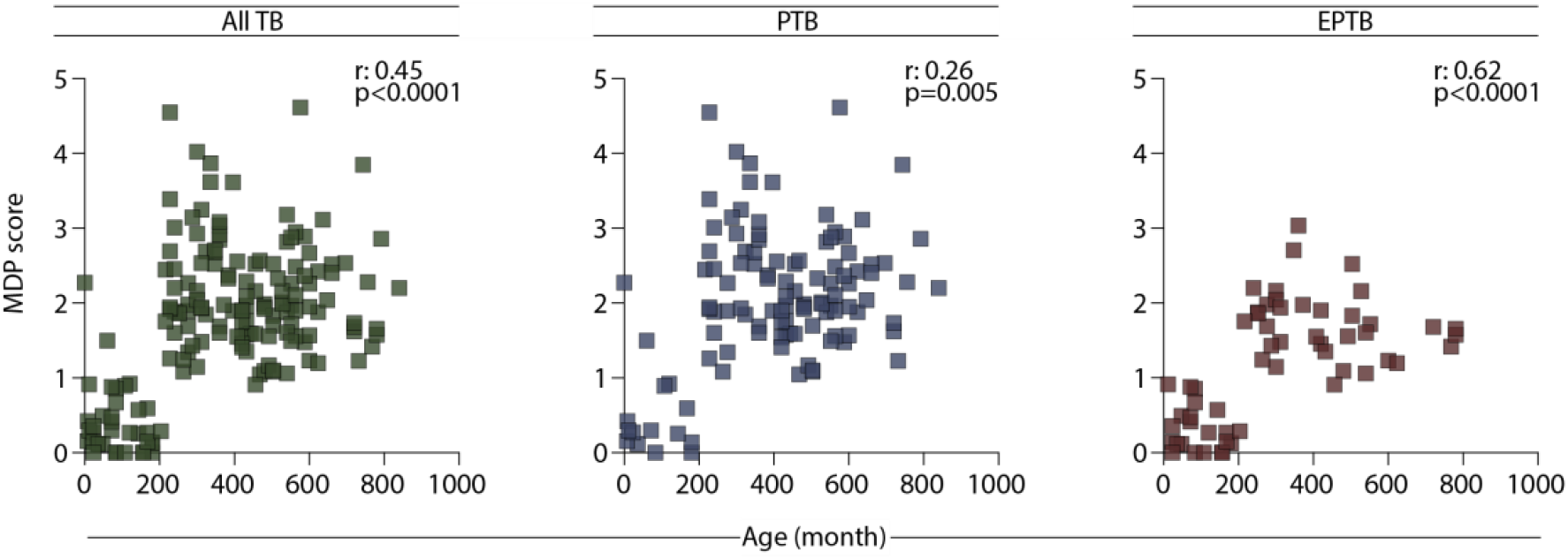
Association between Age and MDP score in patients with tuberculosis. Correlation between Age and MDP score was assessed using the Spearman rank test in the different study groups.

## References

1. Organization WH. Global Tuberculosis Report. 2019.

2. Pai M, Behr MA, Dowdy D, et al. Tuberculosis. Nat Rev Dis Primers 2016; 2:16076.

3. Starke JR. Resurgence of tuberculosis in children. Pediatr Pulmonol Suppl 1995; 11:16–7.

4. Schepisi MS, Motta I, Dore S, Costa C, Sotgiu G, Girardi E. Tuberculosis transmission among children and adolescents in schools and other congregate settings: a systematic review. New Microbiol 2019; 41:282–90.

5. Organization WH. A Research Agenda for Childhood Tuberculosis. 2007.

6. Li J, Sun L, Xu F, et al. Characterization of plasma proteins in children of different Mycobacterium tuberculosis infection status using label-free quantitative proteomics. Oncotarget 2017; 8:103290–301.

7. Silveira-Mattos PS, Barreto-Duarte B, Vasconcelos B, et al. Differential expression of activation markers by Mycobacterium tuberculosis-specific CD4+ T-cell distinguishes extrapulmonary from pulmonary tuberculosis and latent infection. Clin Infect Dis 2019.

8. Oliveira-de-Souza D, Vinhaes CL, Arriaga MB, et al. Molecular degree of perturbation of plasma inflammatory markers associated with tuberculosis reveals distinct disease profiles between Indian and Chinese populations. Sci Rep 2019; 9:8002.

9. Vinhaes CL, Oliveira-de-Souza D, Silveira-Mattos PS, et al. Changes in inflammatory protein and lipid mediator profiles persist after antitubercular treatment of pulmonary and extrapulmonary tuberculosis: A prospective cohort study. Cytokine 2019; 123:154759.

10. Andrade BB, Pavan Kumar N, Amaral EP, et al. Heme Oxygenase-1 Regulation of Matrix Metalloproteinase-1 Expression Underlies Distinct Disease Profiles in Tuberculosis. J Immunol 2015; 195:2763–73.

11. Albuquerque VVS, Kumar NP, Fukutani KF, et al. Plasma levels of C-reactive protein, matrix metalloproteinase-7 and lipopolysaccharide-binding protein distinguish active pulmonary or extrapulmonary tuberculosis from uninfected controls in children. Cytokine 2019; 123:154773.

12. Pavan Kumar N, Anuradha R, Andrade BB, et al. Circulating biomarkers of pulmonary and extrapulmonary tuberculosis in children. Clin Vaccine Immunol 2013; 20:704–11.

13. Blondel VD, Guillaueme, J., Lamblotte, R. & Lefebvre, E. Fast unfolding of communities in large networks. Journal of Statistical Mechanics: Theory and Experiment 2008.

14. S. BMH, M. J. In International AAAI Conference on Weblogs and Social Media, 2009.

15. Mayer-Barber KD, Andrade BB, Oland SD, et al. Host-directed therapy of tuberculosis based on interleukin-1 and type I interferon crosstalk. Nature 2014; 511:99–103.

16. Johnson WE, Li C, Rabinovic A. Adjusting batch effects in microarray expression data using empirical Bayes methods. Biostatistics 2007; 8:118–27.

17. Manabe YC, Andrade BB, Gupte N, et al. A Parsimonious Host Inflammatory Biomarker Signature Predicts Incident TB and Mortality in Advanced HIV. Clin Infect Dis 2019.

18. Rousu J, Agranoff DD, Sodeinde O, Shawe-Taylor J, Fernandez-Reyes D. Biomarker discovery by sparse canonical correlation analysis of complex clinical phenotypes of tuberculosis and malaria. PLoS Comput Biol 2013; 9:e1003018.

19. Pankla R, Buddhisa S, Berry M, et al. Genomic transcriptional profiling identifies a candidate blood biomarker signature for the diagnosis of septicemic melioidosis. Genome Biol 2009; 10:R127.

20. Prada-Medina CA, Fukutani KF, Pavan Kumar N, et al. Systems Immunology of Diabetes-Tuberculosis Comorbidity Reveals Signatures of Disease Complications. Sci Rep 2017; 7:1999.

21. Amaral EP, Ribeiro SC, Lanes VR, et al. Pulmonary infection with hypervirulent Mycobacteria reveals a crucial role for the P2X7 receptor in aggressive forms of tuberculosis. PLoS Pathog 2014; 10:e1004188.

22. Martinez L, Zar HJ. Tuberculin conversion and tuberculosis disease in infants and young children from the Drakenstein Child Health Study: A call to action. S Afr Med J 2018; 108:247–8.

23. Sigal GB, Segal MR, Mathew A, et al. Biomarkers of Tuberculosis Severity and Treatment Effect: A Directed Screen of 70 Host Markers in a Randomized Clinical Trial. EBioMedicine 2017; 25:112–21.

24. Scriba TJ, Coussens AK, Fletcher HA. Human Immunology of Tuberculosis. Microbiol Spectr 2016; 4.

25. Cliff JM, Kaufmann SH, McShane H, van Helden P, O'Garra A. The human immune response to tuberculosis and its treatment: a view from the blood. Immunol Rev 2015; 264:88–102.

26. Mayer-Barber KD, Sher A. Cytokine and lipid mediator networks in tuberculosis. Immunol Rev 2015; 264:264–75.

27. Mayer-Barber KD, Yan B. Clash of the Cytokine Titans: counter-regulation of interleukin-1 and type I interferon-mediated inflammatory responses. Cell Mol Immunol 2017; 14:22–35.

28. Novikov A, Cardone M, Thompson R, et al. Mycobacterium tuberculosis triggers host type I IFN signaling to regulate IL-1beta production in human macrophages. J Immunol 2011; 187:2540–7.

29. Mayer-Barber KD, Andrade BB, Barber DL, et al. Innate and adaptive interferons suppress IL-1alpha and IL-1beta production by distinct pulmonary myeloid subsets during Mycobacterium tuberculosis infection. Immunity 2011; 35:1023–34.

30. Fredenburgh LE, Perrella MA, Mitsialis SA. The role of heme oxygenase-1 in pulmonary disease. Am J Respir Cell Mol Biol 2007; 36:158–65.

31. Origassa CS, Camara NO. Cytoprotective role of heme oxygenase-1 and heme degradation derived end products in liver injury. World J Hepatol 2013; 5:541–9.

32. Andrade BB, Singh A, Narendran G, et al. Mycobacterial antigen driven activation of CD14++CD16− monocytes is a predictor of tuberculosis-associated immune reconstitution inflammatory syndrome. PLoS Pathog 2014; 10:e1004433.

33. Hsu DC, Breglio KF, Pei L, et al. Emergence of Polyfunctional Cytotoxic CD4+ T Cells in Mycobacterium avium Immune Reconstitution Inflammatory Syndrome in Human Immunodeficiency Virus-Infected Patients. Clin Infect Dis 2018; 67:437–46.

